# A Human Angelman Syndrome Class II Pluripotent Stem Cell line with Fluorescent Paternal *UBE3A* Reporter

**DOI:** 10.1101/2025.07.12.664539

**Authors:** Gautami R. Kelkar, Samantha R. Stuppy, Dilara Sen, Z. Begum Yagci, Linna Han, Lexi Land, Jessica K. Hartman, Albert J. Keung

## Abstract

**Introduction:** Angelman Syndrome (AS) is characterized in large part by the loss of functional UBE3A protein in mature neurons. A majority of AS etiologies is linked to deletion of the maternal copy of the *UBE3A* gene and epigenetic silencing of the paternal copy. A common therapeutic strategy is to unsilence the intact paternal copy thereby restoring UBE3A levels. Identifying novel therapies has been aided by a *UBE3A-YFP* reporter mouse model. This study presents an analogous fluorescent *UBE3A* reporter system in human cells.

**Methods:** Previously derived induced Pluripotent Stem Cells (iPSCs) with a Class II large deletion at the UBE3A locus are used in this study. *mGL* and *eGFP* are integrated downstream of the endogenous *UBE3A* using CRISPR/Cas9. These reporter iPSCs are differentiated into 2D and 3D neural cultures to monitor long-term neuronal maturation. Green fluorescence dynamics are analyzed by immunostaining and flow cytometry.

**Results:** The reporter is successfully integrated into the genome and reports paternal *UBE3A* expression. Fluorescence expression gradually reduces with *UBE3A* silencing in neurons as they mature. Expression patterns also reflect expected responses to molecules known to reactivate paternal *UBE3A*.

**Discussion:** This human-cell-based model can be used to screen novel therapeutic candidates, facilitate tracking of *UBE3A* expression in time and space, and study human-specific responses. However, its ability to restore UBE3A function cannot be studied using this model. Further research in human cells is needed to engineer systems with functional UBE3A to fully capture the therapeutic capabilities of novel candidates.

## 1 Introduction

*UBE3A* encodes an E3 ubiquitin ligase (UBE3A, E6-AP) that is integral for several cellular processes that govern cortical development (LaSalle, Reiter, and Chamberlain 2015; Khatri and Man 2019). While it is biallelically expressed in most early developmental cell types, *UBE3A* switches to monoallelic expression in mature neurons (Albrecht et al. 1997; Rougeulle, Glatt, and Lalande 1997; Vu and Hoffman 1997). The paternal allele is epigenetically silenced by a long non-coding RNA transcript, *UBE3A-ATS*, leaving only the maternal allele actively transcribed. Loss or loss of function of the maternal allele leads to a dearth of UBE3A and is the primary linkage to the neurodevelopmental disorder, Angelman Syndrome (AS) (Angelman 1965; Kishino, Lalande, and Wagstaff 1997; Matsuura et al. 1997; Sutcliffe et al. 1997).

The predominant etiologies affecting maternal *UBE3A* are large genomic deletions comprising approximately 70% of all AS (Keute et al. 2021). Since the paternal *UBE3A* allele is still intact, therapeutic strategies often target its unsilencing via inhibition or knockdown of *UBE3A-ATS*. Thus, the ability to track the transcriptional activity of paternal *UBE3A* would facilitate therapeutic screening. It would also benefit the spatiotemporal mapping of allele-specific *UBE3A* expression, which is a highly dynamic and cell-type-specific process (Burette et al. 2018; 2017; Estridge et al. 2025; Judson et al. 2014; Sen, Drobna, and Keung 2021; Sen et al. 2020). The *UBE3A-YFP* reporter mouse model (Dindot et al. 2008) has been leveraged to screen for therapeutic molecules, identifying topoisomerase inhibitors (Huang et al. 2012; H.-M. Lee et al. 2018), anti-sense oligonucleotides (ASOs) (Clarke et al. 2024; D. Lee et al. 2023; Meng et al. 2015), CRISPR-Cas9 gene therapies (Bazick et al. 2024; Schmid et al. 2021; Wolter et al. 2020), and small molecules like PHA533533 (Vihma et al. 2024). Future screens, preclinical validations, and research studies would all benefit from an analogous human system (Sen, Drobna, and Keung 2021). It would account for species-specific differences in (epi)genetic variation, developmental timescales, dose-response characteristics, and toxicities (Chen 2016). In addition, such models could benefit mechanistic studies on *UBE3A* dynamics and its role in cellular processes (Condon et al. 2013; Filonova et al. 2014; Grier, Carson, and Lagrange 2015; Hillman et al. 2017; Jones et al. 2016; Judson et al. 2014; Kaphzan et al. 2012; McCoy et al. 2017; Sonzogni et al. 2020). In a few cases, compound hits from the mouse model were then validated on human iPSC-derived neurons, but these cellular systems required the use of antibody labeling or amplification which need laborious fixation steps and have known challenges of sensitivity and specificity (Dindot et al. 2023; Sen et al. 2020).

Here, we modify a human AS Class II patient-derived induced pluripotent stem cell (iPSC) line by knocking in a fluorescent reporter in frame with paternal *UBE3A* (Chamberlain et al. 2010). These iPSCs are differentiated into 2D neurons as well as cerebral organoids to observe changes in fluorescence levels with time, in specific cell types, and with neural cell maturation (Ciceri et al. 2024; Lancaster et al. 2013). We also assess *UBE3A* reactivation upon exposure to topoisomerase inhibitors.

## 2 Materials and Methods

### 2.1 DNA plasmids

Two donor plasmids expressing GSLinker-eGFP and 2A-eGFP were generated. However, the fluorescence signal was not successfully detected in iPSCs. Hence, the final *UBE3A*-targeting construct was designed for both direct and indirect expression: UBE3A-GSLinker-mGreenLantern-IRES-NLS-eGFP. First, GSLinker-eGFP (pDS44-GSLinker) was subcloned into the CIRTS-1: ORF5-TBP6.7-Pin domain-NLS plasmid (Addgene #132543) using restriction digestion at NdeI and MfeI, followed by T4 ligation. The resulting plasmid (pSS12-GSLinker-eGFP) was cut using BseRI and BstEII to remove the GS linker and to perform Gibson assembly to add PAM mutations and an IRES-NLS sequence. Finally, a GSLinker-mGreenLantern fragment was added before the IRES via Gibson assembly to obtain pSS23-GSLinker-mGreenLantern-IRES-NLS-eGFP. For the Cas9-gRNA construct, pX330-U6-Chimeric_BB-Cbh-hSpCas9 (Addgene #42230) was used to clone the gRNA (AGGCCATCACGTATGCCAA (C. Sirois 2018)). The donor construct pSS22-IRES-NLS-eGFP along with information about the GSLinker-mGreenLantern fragment (# 241837) and Cas9-gRNA construct pDS48 (# 241839) used in the present study are available from Addgene.

### 2.2 Cell transfection and genomic PCR screening

AS Class II deletion iPSCs (developed by Chamberlain and colleagues, and procured from Kerafast) were transfected via electroporation (Chamberlain et al. 2010). The experiment used 1.5 μg of pSS23 and 1.5 μg of pDS48 in a 10 μl Neon^™^ Transfection System (Invitrogen) reaction containing 50,000 cells. The following conditions were used: 950 Volts - 2 pulse - 30 width. Cells were immediately seeded post-transfection onto a growth factor-reduced Matrigel-coated (Corning) CellRaft AIR^®^ array from Cell Microsystems and allowed to recover for about a week with gentle mTeSR Plus (StemCell Technologies) media changes. Upon expansion, the cells were dissociated with Accutase (BioLegend) and seeded into 96-well plates (Corning COSTAR^™^). Replicate plates were generated and screened by PCR of genomic DNA. All media contained CloneR (StemCell Technologies) per the manufacturer’s instructions from transfection through expansion and re-plating. Genomic PCR screens were performed with primer sets SRS21 + SRS24, followed by Sanger sequencing to identify a polyclonal population. These cells were further sorted using the CellRaft AIR^®^ system, and the aforementioned PCR screening was repeated. An additional PCR screening step was performed with SRS21 + DSP206 primers, followed by Nanopore sequencing (performed by SNPsaurus) to identify 3 monoclonal populations of reporter iPSCs. The sequences for primers can be found in Supplementary Table S1.

### 2.3 Human Pluripotent Stem Cell (hPSC) culture

The parental AS Class II deletion iPSCs, the reporter iPSCs, and the H9_*UBE3A m-/p-*_ ESCs (from Dr. Stormy Chamberlain) were maintained on growth factor-reduced Matrigel (Corning) coated 6-well plates (Corning COSTAR^™^, Fisher Scientific) in mTeSR Plus (StemCell Technologies). Cells were passaged every 3-5 days as necessary using 0.5 mM EDTA (Invitrogen).

### 2.4 RNA extraction and qPCR

Parental and reporter iPSCs were washed with cold 1X PBS (Gibco) and total RNA was isolated using the RNeasy^®^ Mini Kit (Qiagen) following manufacturer’s instructions. RNA samples were treated with the Turbo DNA-free^™^ kit (Invitrogen) to remove DNA contamination. For qPCR, cDNA synthesis was performed using the iScript Advanced cDNA Synthesis Kit (BIO-RAD) according to the manufacturer’s protocol. Primers SRS35 and SRS36 were used to amplify 2 regions: one within mGL and eGFP each, and primers SRS37 and SRS38 were used to amplify a region within eGFP only. GAPDH was used as the reference gene. Data was analyzed in MS Excel and is presented as ΔCt relative to GAPDH. The sequences for primers can be found in Supplementary Table S1.

### 2.5 2D neural culture

All media for neural cultures were sourced from StemCell Technologies, and the procedures adhered to their established protocols. iPSCs were differentiated to Neural Progenitor Cells (NPCs) using the StemDiff^™^ SMADi neural induction kit and stored in liquid nitrogen. Frozen NPCs were thawed and grown to confluency before being differentiated into neural precursor cells. Confluent NPCs were passed using Accutase (BioLegend) to a well of a 6-well plate (Corning COSTAR^™^, Fisher Scientific) at 1.3×10^5^ cells/cm^2^ using the StemDiff^™^ Forebrain Neuron Differentiation Kit. Upon reaching confluency, these cells were passed using Accutase into 8-well chamber slides (Falcon) at 2.00×10^4^ cells/cm^2^ and maintained using the StemDiff^™^ Forebrain Neuron Maturation Kit. The media was changed every 2-3 days. To accelerate neuron maturation, 4 μM GSK343 (MilliporeSigma) was added at every media change (Ciceri et al. 2024).

### 2.6 Cerebral organoid culture

Whole-brain organoids were generated from AS Class II deletion parental and reporter stem cell lines. hiPSCs were allowed to reach 80-90% confluency before they were dissociated into a single-cell suspension using Accutase (BioLegend) and re-plated in low-adhesion U-bottom 96-well plates (Corning COSTAR^™^, Fisher Scientific) at 12,000-15,000 cells/well in organoid generation media. Organoids were generated and maintained by modifying the protocol in (Lancaster et al. 2013). To accelerate neuron maturation, 4 μM GSK343 (MilliporeSigma) was added to the organoids between Days 17-25, with partial media change carried out every other day (Ciceri et al. 2024).

### 2.7 Histology and immunofluorescence

Tissues were fixed in 4% paraformaldehyde for 20 minutes at 4 ^○^C followed by washing in 1X PBS (Gibco) three times for 10 minutes. Tissues were allowed to equilibrate in 30% sucrose overnight and then embedded in 10% / 7.5% gelatin/sucrose. Embedded tissues were frozen in an isopentane and dry ice bath at -30 to -50 ^○^C and stored at -80 ^○^C. Prior to analysis, they were cryosectioned into 30 μm slices using a cryoStat (ThermoFisher). iPSCs and 2D neural cultures plated in 8-well chamber slides (Falcon) were fixed in 4% paraformaldehyde for 10 minutes at room temperature followed by washing in 1X PBS (Gibco) three times for 5 minutes. For immunohistochemistry, organoid sections were blocked and permeabilized in 1% Triton X-100 and 5% normal donkey serum (Jackson Immunoresearch Labs) in 1X PBS. For 2D cultures, 1% normal donkey serum in 1% Triton X-100 was used instead. Both 2D cultures and organoid sections were then incubated with primary antibodies in 10% Triton X-100, 1% normal donkey serum in UltraPure^™^ water (Invitrogen) and 10X PBS overnight at 4 ^○^C at the following dilutions: UBE3A (rabbit, Bethyl Laboratories, 1:250 or mouse, MilliporeSigma, 1:1000 (Figure 2A)), GFP (chicken, abcam, 1:2000 for iPSCs and 2D neural cultures; 1:3000 for organoid sections), OCT4 (rabbit, Cell Signaling, 1:200), SOX2 (goat, R&D Systems, 1:200), and TUJ1 (mouse, MilliporeSigma, 1:500). Following primary antibodies, sections were incubated with secondary antibodies - donkey Alexa Fluor 488, 546, and 647 conjugates (Invitrogen, 1:250) in 10% Triton X-100, 1% normal donkey serum in UltraPure^™^ water (Invitrogen) and 10X PBS for 2 hours at room temperature, and the nuclei were stained with DAPI (Invitrogen). Slides were mounted using ProLong^™^ Antifade Diamond (Invitrogen). Images were taken with Nikon A1R confocal microscope (Nikon Instruments).

**Figure 1.**
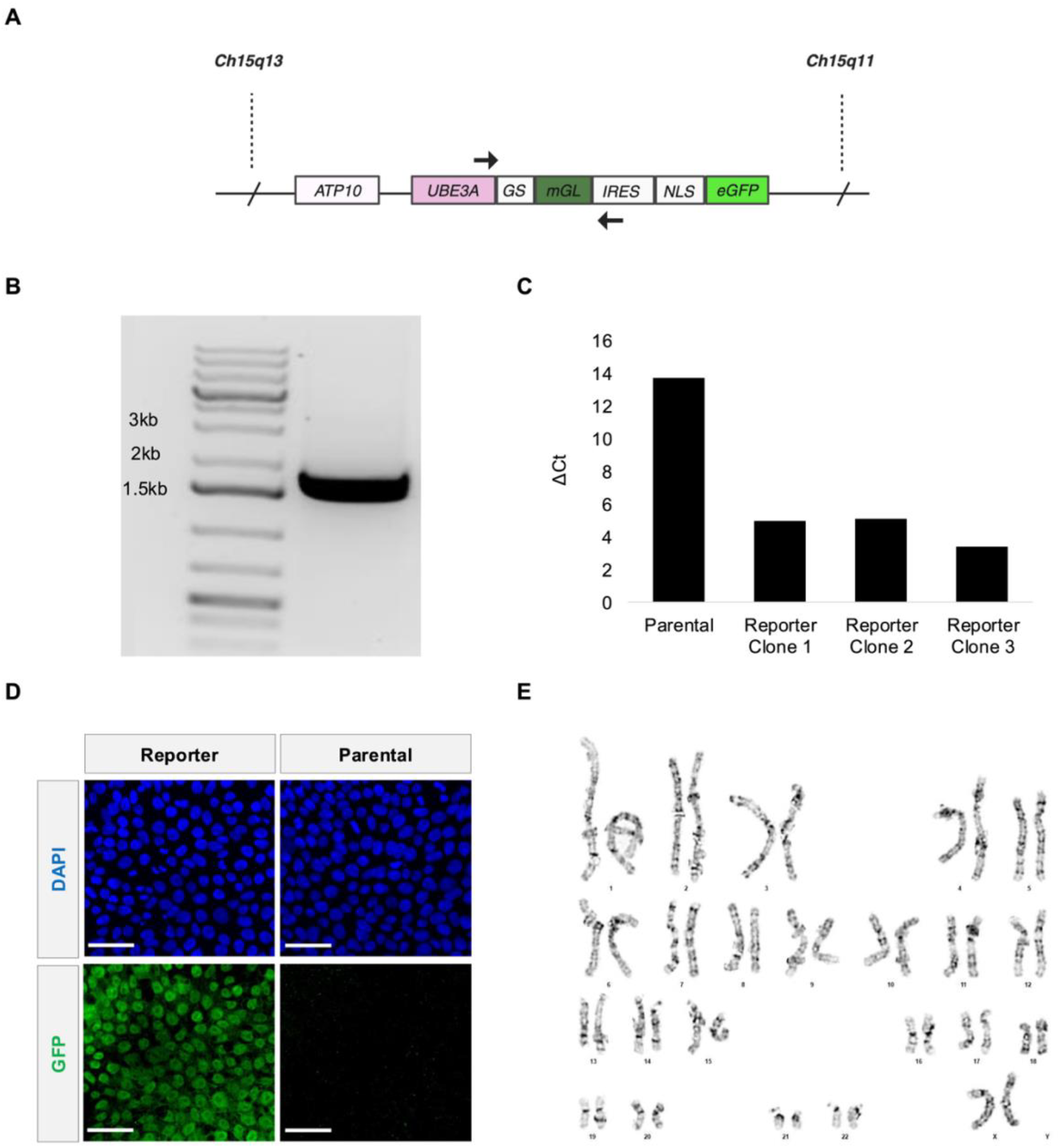
Fluorescent *UBE3A* reporter knocked into AS Class II deletion iPSCs. A) Illustration of the *UBE3A* locus in paternal Chromosome 15 after reporter integration. B) Primers SRS21 (forward) and SRS24 (reverse) targeting the *UBE3A*-reporter region shown in 1A amplified a band that is ∼ 1.756kb from genomic DNA of the polyclonal reporter cells. C) qRT-PCR measurements of fluorescent reporter mRNA (Supplementary Figure 1A). ΔCt values relative to GAPDH presented for the parental iPSCs and the three monoclonal reporter iPSCs. D) Representative confocal microscopy images showing GFP expression in the nuclei (DAPI) of AS Class II deletion parental iPSCs (right) and the edited reporter iPSCs (left). Scale bars are 50 µm. E) G-banded chromosomes show normal chromosome counts of 46 XX. The karyotype analysis was performed by LabCorp.

**Figure 2.**
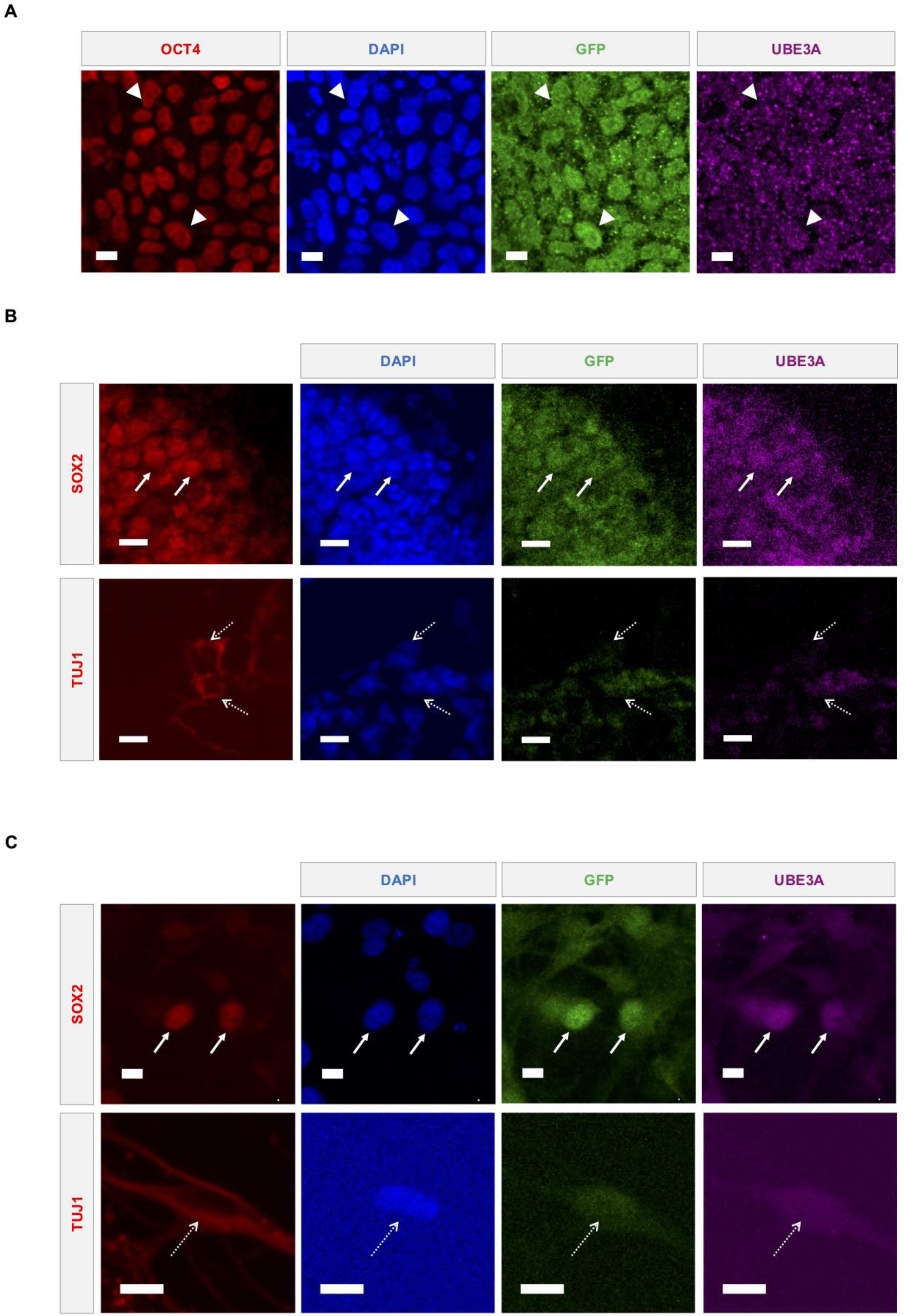
Direct UBE3A labeling and reporter fluorescence coexpress in multiple cell types. A) Representative confocal microscopy images showing UBE3A and GFP expression in the nuclei (DAPI) of reporter iPSCs. OCT4 was used as a marker to identify iPSCs. Representative confocal microscopy images showing UBE3A and GFP expression in the nuclei (DAPI) of cells in edited reporter iPSC-derived B) cerebral organoids and C) 2D neural cultures. SOX2+ neural precursors shown using solid arrows and TUJ1+ neurons marked by dashed arrows. All scale bars are 10 µm.

### 2.8 Image analysis and quantification

All samples within experiments were imaged using the same microscope settings, and intensities for all channels were adjusted identically in FIJI (Schindelin et al. 2012). For Figure 3, 3 organoids per time-point, and 3 regions per organoid were imaged. For UBE3A and GFP intensity measurements over time, 20 cells of one type were analyzed in each image (180 cells per condition). Intensities for each channel within each image were normalized to a range of 0-1 as follows.

**Figure 3.**
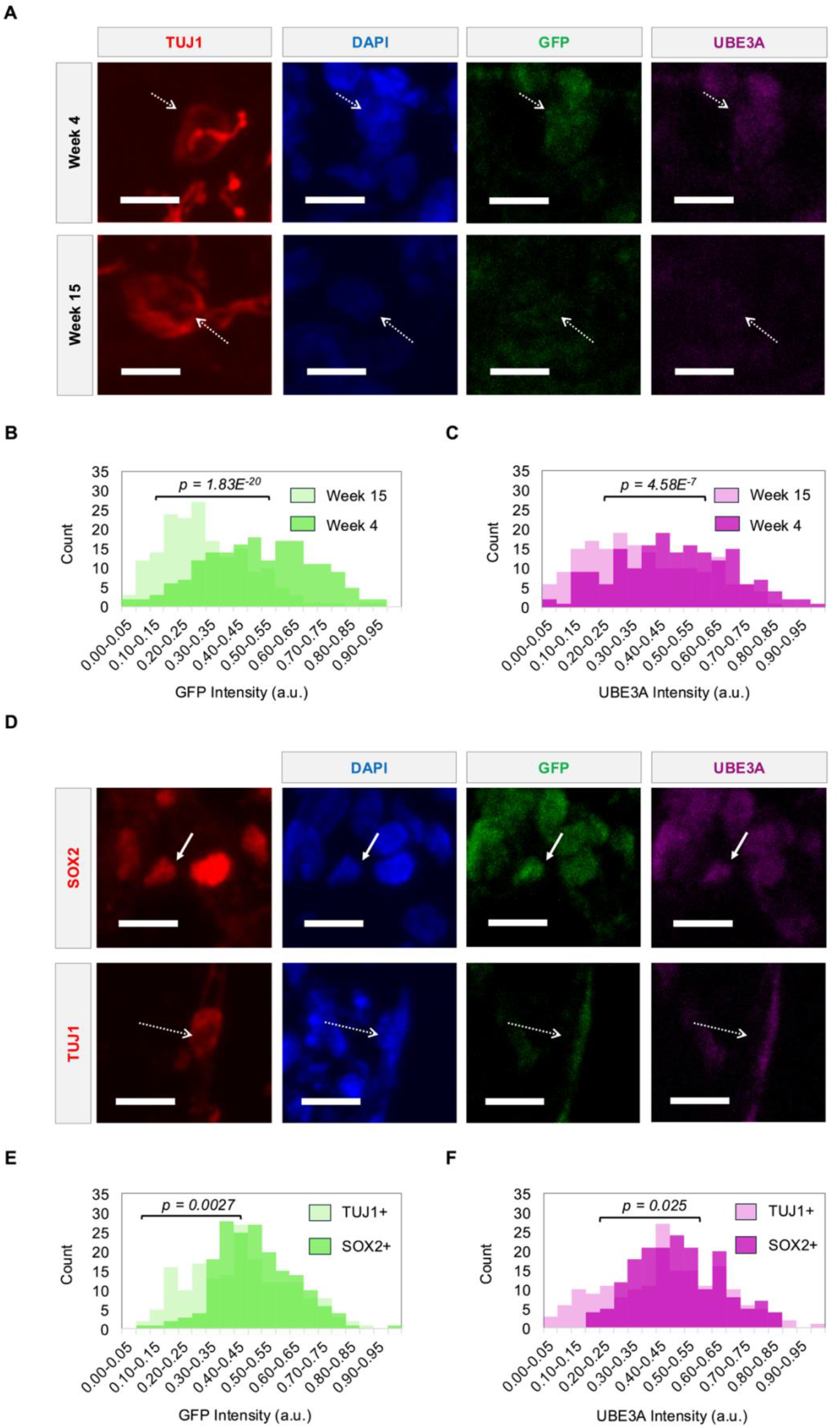
Reporter fluorescence tracks UBE3A expression dynamics during neural differentiation. A) Representative images showing UBE3A and GFP expression in the nuclei (DAPI) of TUJ1+ neurons (dashed arrows) in cerebral organoids at week 4 and week 15. B) GFP and C) UBE3A intensities of nuclei in TUJ1+ neurons at weeks 4 and 15. D) Representative images showing UBE3A and GFP expression in the nuclei of SOX2+ precursors (solid arrows) and TUJ1+ neurons (dashed arrows) in cerebral organoids at week 9. E) GFP and F) UBE3A intensities of nuclei in TUJ1+ neurons and SOX2+ precursors at week 9. Populations were compared using a two-sample t-test, n = 180 cells (3 organoids). All scale bars are 10 µm. a.u. - arbitrary units.

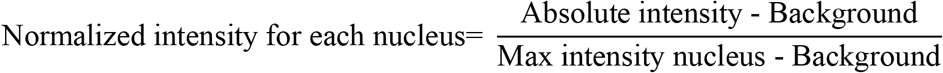

### 2.9 Topoisomerase inhibitor treatment

Topotecan hydrochloride and irinotecan hydrochloride (both from Molcan Corporation) were directly added to 17-week-old AS reporter organoids at a final concentration of 1μM in culture. 0.2% DMSO in water was used as the vehicle control. Organoids were incubated at 37 ^○^C on an orbital shaker at 90 RPM for 72 hours before being harvested for analysis.

### 2.10 Flow cytometry and data visualization

Adherent iPSCs were washed with 1mL 1X PBS (Gibco), incubated with Accutase (BioLegend) for 5 minutes at 37 ^○^C, and dissociated into a single-cell suspension. Organoids were also washed with 1X PBS, incubated in Accutase for 3 rounds of 10 minutes each at 37 ^○^C with pipetting to break up the organoids between rounds, ultimately obtaining a single-cell suspension. For both iPSCs and organoids, the single-cell suspensions were centrifuged at 300 g for 5 minutes, Accutase was removed, and the cells were re-suspended in 300 nM DAPI (Invitrogen) in 1X PBS. The samples were then run on the flow cytometer (MACSQuant^®^ VYB) with identical volumes for all samples within an experiment. The data was processed and visualized using FlowJo^™^ v10.10.0 software (BD Life Sciences).

### 2.11 UBE3A ubiquitin ligase activity assay

The assay followed the protocol described in (Han, Yagci, and Keung 2025). Briefly, lysates from all three cell lines (3 biological replicates per cell line) were collected using the Pierce^™^ IP lysis buffer (Fisher Scientific). The total amount of protein present in the cell lysates was measured using Pierce^™^ BCA Protein Assay Kit (ThermoFisher Scientific) following which the lysates were concentrated using Amicon^®^ ultra centrifugal filters (MilliporeSigma, 50kDa MWCO, 0.5ml). The concentrated lysate was used in the ubiquitin ligase activity reaction mixture. The reaction components and their approximate working concentrations/amounts were as follows: 10X reaction buffer (10 mM), Mg^2+^/ATP (10 mM, Enzo Life Sciences), UBE1 (100 nM, R&D Systems), UBE2 (1 µM, R&D Systems), HPV-E6 (1.5 µM, R&D Systems), custom p53 substrate (5.4 µg, Addgene plasmid # 233738) and ubiquitin-fluorescein (10 µM, R&D Systems). Amicon^®^ filter-concentrated IP lysis buffer was used as the blank control. The samples were incubated at 37 ^○^C for an hour. HisPur^™^ Ni-NTA magnetic beads (Fisher) were used to pull down p53 substrates. To calculate the normalized fluorescence readings, the fluorescence reading of the IP lysis buffer control was first subtracted from the readings of the samples, and then these readings were normalized to their corresponding approximate protein amounts.

## 3 Results

### 3.1 Fluorescent *UBE3A* reporter knocked into AS Class II deletion iPSCs

Prior correspondence with researchers in the field working in both mouse and human systems noted issues detecting fluorescence signals with direct fusions of fluorescence proteins to UBE3A, potentially due to the misfolding of the fluorescent protein. This therefore necessitates the use of fixation and antibody labeling of the fluorescent protein (Dindot et al. 2008; H.-M. Lee et al. 2018; Huang et al. 2012; Meng et al. 2015; D. Lee et al. 2023; Clarke et al. 2024; Schmid et al. 2021; Vihma et al. 2024; Judson et al. 2014; McCoy et al. 2017; Jones et al. 2016; Condon et al. 2013; Hillman et al. 2017). In addition, direct fusions to UBE3A in human cell lines, specifically, seemed to face issues of detection even with antibody labeling (Chen 2016). Given these potential challenges, here we edit an AS Class II iPSC line harboring a deletion of 15q11-q13 on the maternal chromosome (Chamberlain et al. 2010), with three different constructs. All three constructs target the c-terminus of paternal *UBE3A* using CRISPR/Cas9-induced homology directed repair. These include in frame insertions prior to the *UBE3A* stop codon of: *GSLinker-eGFP*; *2A-eGFP*; and *GSLinker-mGreenLantern (mGL)-IRES-NLS-eGFP*. The third construct is designed for two reasons. First, it may maximize the signal possible from the construct using both a brighter mGL fluorophore (Campbell et al. 2020) and two fluorophores combined. Second, eGFP being translated as a separate protein increases the chance of detecting gene activation without the need for antibody labeling and avoiding potential disruption to UBE3A and fluorophore folding. Interestingly, only the third construct (Figure 1A) yields detectable signal. Genotyping indicates this construct is successfully integrated into the genome (Figure 1B). The Cell Microsystems’ CellRaft AIR^®^ system enables isogenic clonal populations to be further isolated. Expression of the reporter transcript is confirmed using qRT-PCR (Figure 1C; Supplementary Figure S1) for all three clonal populations. One clone is selected for further analysis. At the protein level, the reporter iPSCs show higher antibody-enhanced green fluorescence compared to the parental iPSCs in confocal imaging (Figure 1D). The reporter cell line maintains a normal karyotype (Figure 1E; Supplementary Figure S2).

### 3.2 Direct UBE3A labeling and reporter fluorescence coexpress in multiple cell types

The coexpression of fluorescent reporters with UBE3A is essential for their utility. Antibody-enhanced fluorescence imaging indicates coexpression of the fluorescent proteins with UBE3A (Figure 2A), albeit with the UBE3A signal being relatively noisier than the reporter. In AS deletion etiologies, paternal *UBE3A* is known to be expressed in pluripotent stem cells, neural precursors, and immature neurons of murine (Judson et al. 2014) and human (Chamberlain et al. 2010; Sen et al. 2020) models. As expected based on this prior literature, coexpression is also observed in SOX2+ neural precursors and TUJ1+ neurons of 4-week-old cerebral organoids generated from the reporter iPSCs (Figure 2B). Similarly, in 9-week-old 2D neural cultures, expression is observed in both SOX2+ and TUJ1+ cells (Figure 2C). The nuclear localization of UBE3A is consistent with previous reports studying neurodevelopmental cell types in human iPSC-derived models for AS (Zampeta et al. 2020; Munshi 2019).

### 3.3 Reporter fluorescence tracks UBE3A expression dynamics during neural differentiation

During neurodevelopment, paternal *UBE3A* is gradually silenced over time during neuronal differentiation and maturation (Stanurova et al. 2016; Hsiao et al. 2019). Fluorescence imaging of long-term cerebral organoid cultures at weeks 4, 9, 12, and 15 show that the reporter tracks this silencing. Consistent with prior findings, UBE3A exhibits strong nuclear localization up to week 12, which becomes weaker and more diffuse by week 15 (Sen et al. 2020) (Figure 3A). Reporter expression mirrors this pattern. Quantitative analysis reveals a significant decline in nuclear fluorescence intensities of both UBE3A and the reporter by week 12 (Supplementary Figure S3), and this is sustained through week 15 (Figures 3B and 3C). Furthermore, silencing dynamics also track cellular differentiation where TUJ1+ neurons consistently show weaker nuclear UBE3A and reporter signals compared to their signals within SOX2+ neural precursors (Figure 3D). Notably, this difference is evident as early as week 9, with neurons displaying significantly lower intensities than precursors (Figures 3E and 3F; Supplementary Figure S4). These results confirm the silencing of paternal *UBE3A* during neural maturation and demonstrate that the reporter faithfully tracks this dynamic expression.

### 3.4 Unamplified native reporter fluorescence is detectable by flow cytometry

For both the *UBE3A-YFP* mouse model (Dindot et al. 2008) and the imaging experiments in this study, antibody-based signal enhancement is required to visualize fluorescent protein expression. Flow cytometry provides a potential alternative measurement method with high sensitivity as well as single-cell resolution. Indeed, we observe that flow cytometry is able to distinguish unedited parental cells from cells edited with the reporter (Figure 4). Furthermore, 17-week-old cerebral organoids display intermediate fluorescence levels above the parental iPSC baseline and below the reporter iPSC signal (Figure 4). This reflects a decline in reporter fluorescence over time, consistent with *UBE3A* silencing. These results demonstrate that flow cytometry can reliably detect endogenous reporter fluorescence and its silencing trajectory, offering an antibody-independent method to monitor UBE3A dynamics.

**Figure 4.**
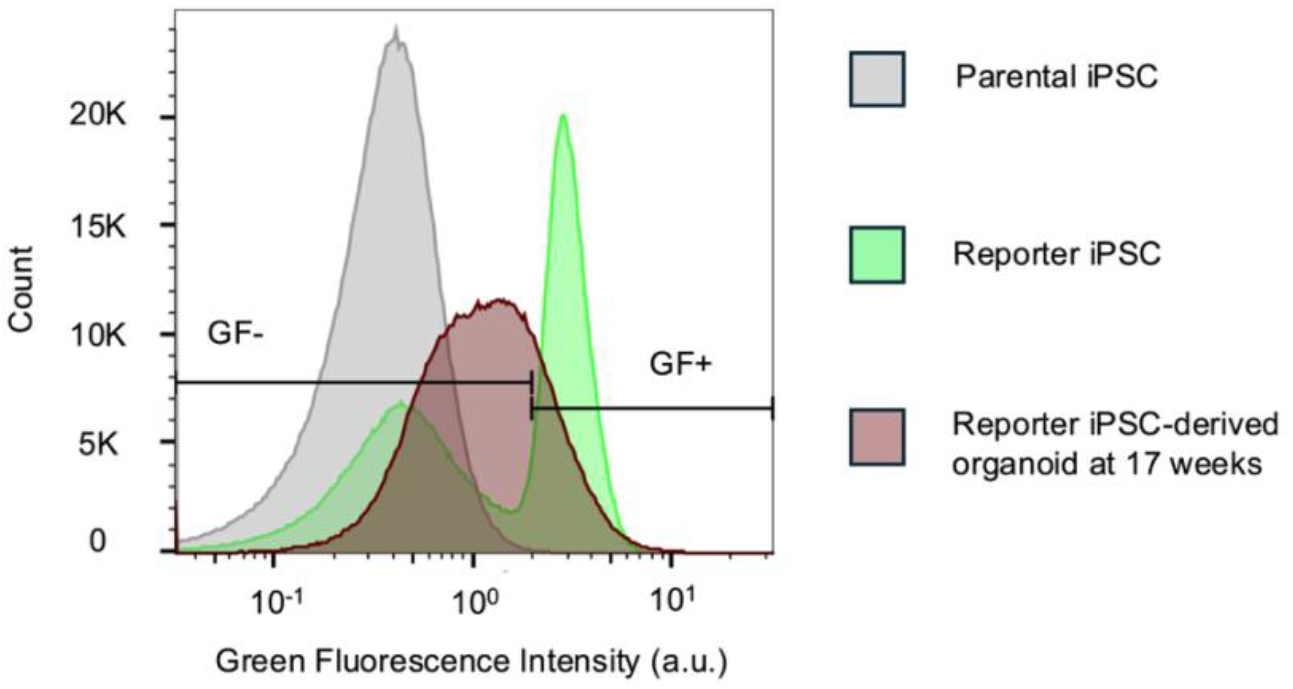
Unamplified native reporter fluorescence is detectable by flow cytometry. Flow cytometry histograms comparing green fluorescence (GF) intensities in parental and reporter iPSCs and dissociated 17-week-old reporter organoids. The iPSCs were used to determine the GF +/- gate. For the organoids, 76.8% cells were GF-. a.u. - arbitrary units.

### 3.5 Reporter fluorescence changes in response to topoisomerase inhibitors

Paternal *UBE3A*, which becomes epigenetically silenced in mature neurons, can be reactivated through various molecular strategies (de Almeida, Tonazzini, and Daniele 2025). A potential application of the reporter cell line is to facilitate screening of novel compounds capable of inducing such reactivation. One of the first classes of molecules discovered to reactivate paternal *UBE3A* is topoisomerase inhibitors, specifically topotecan, which has previously reactivated *UBE3A* in human cerebral organoids (Sen et al. 2020), and irinotecan, which demonstrated similar effects in a *UBE3A-YFP* mouse model. They function by prematurely terminating the *UBE3A-ATS* transcript, thereby permitting expression of *UBE3A* (Huang et al. 2012). The expectation is that topotecan or irinotecan treatment of reporter organoids older than week 15, when both *UBE3A* and reporter fluorescence are largely silenced in neurons, would reactivate reporter fluorescence. We observe an interesting bifurcation of the cell population in 17-week-old organoids following either topotecan or irinotecan treatment. In the bimodal populations, one subpopulation exhibits increased reporter intensity, and the other subpopulation exhibits lower intensities compared to untreated organoids (Figures 5A-C). Flow cytometry gating based on parental (negative control) and reporter (positive control) iPSCs, indicate a higher proportion of reporter-positive cells in organoids treated with topoisomerase inhibitors and also a corresponding increase in mean fluorescence intensity (Supplementary Table S2).

**Figure 5.**
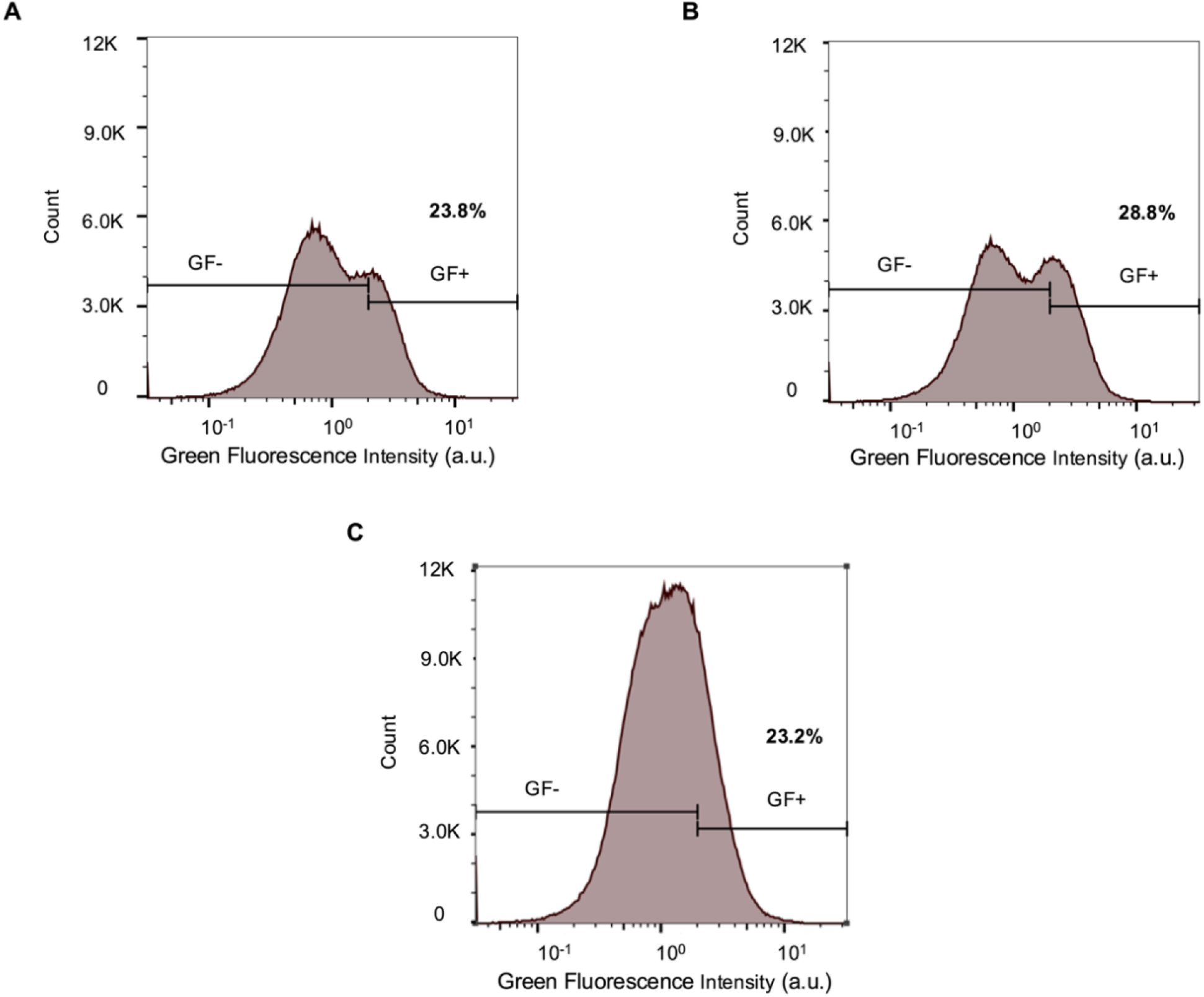
Reporter fluorescence changes in response to topoisomerase inhibitors. Flow cytometry histograms comparing green fluorescence intensities in 17-week-old reporter organoids exposed to A) 1 µM topotecan, B) 1 µM irinotecan, and C) 0.2% DMSO in water (vehicle control). The parental and reporter iPSCs were used to determine the GF +/-gate. a.u. - arbitrary units.

### 3.6 Fusion of the reporter with UBE3A reduces enzyme activity

Fusion of mGL with UBE3A via the GS Linker, while enabling reporting of its transcriptional activation, could affect the enzymatic activity of UBE3A. UBE3A functions as an E3 ligase in the ubiquitin proteasome system (Khatri and Man 2019). To test this, UBE3A ubiquitin ligase activity in the reporter iPSCs can be measured using a recently developed ubiquitin conjugation and pull-down assay (Han, Yagci, and Keung 2025). Using this assay, we observe the ubiquitin ligase activity of the reporter iPSCs is significantly reduced compared to that of the parental iPSCs. H9_*UBE3A m-/p-*_ ESCs, in which *UBE3A* was deleted (C. Sirois 2018; C. L. Sirois et al. 2020; Fink et al. 2017), were used as background control for non-specific activity (Figure 6).

**Figure 6.**
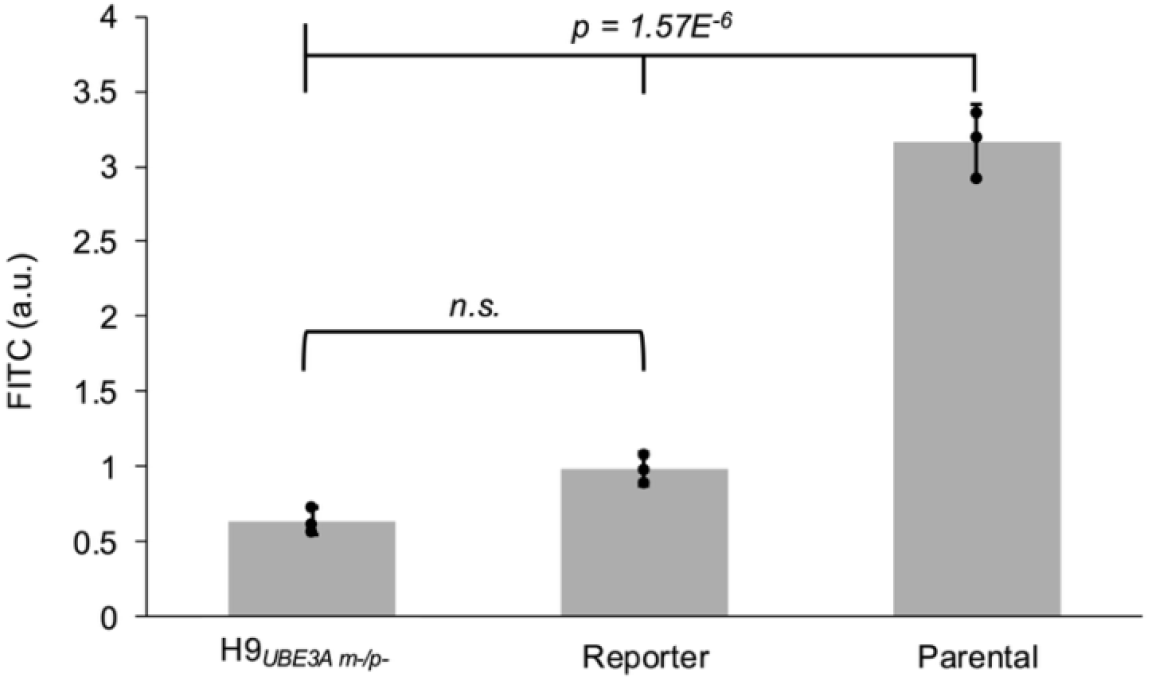
Fusion of the reporter with UBE3A reduces enzyme activity. Normalized fluorescence readings from the UBE3A activity assay performed on the cell lysates of H9_*UBE3A m-/p-*_ ESCs, reporter iPSCs, and parental iPSCs. Error bars represent 95% confidence intervals. Black circles represent each biological replicate (3 per cell line). Full tick mark compared with half tick marks using one-way ANOVA followed by Tukey-Kramer post-hoc test. a.u. - arbitrary units, n.s. - not significant.

## 4 Discussion

The reporter cell line presents both advantages and disadvantages. It could facilitate and accelerate both the exploration of mechanisms regulating *UBE3A* expression as well as the search for new therapeutic compounds. In particular, its compatibility with antibody-free detection using flow cytometry would support rapid screening of novel compound libraries. The single-cell resolution of flow cytometry could also be leveraged to include cell-type-specific antibodies and to query reporter responses in individual cell types. However, a key limitation is that downstream validation of phenotypic recovery in response to potential therapeutics would not be compatible with the reporter cell line as the enzymatic activity of UBE3A is ablated. Instead, the unedited parental cell line along with other AS iPSC lines could be used instead. It is also important to note that the scope of use for this reporter is limited to AS maternal deletion etiologies.

Upon treatment of reporter organoids with topotecan and irinotecan, we observe two subpopulations. One subpopulation exhibits an increase in reporter intensity over the untreated organoids. The number of cells with an increase in intensity was modest, which is important to note as cerebral organoids are inherently heterogeneous. In addition, there was the emergence of a lower intensity subpopulation. Unexpectedly, the mean intensity of this subpopulation is even lower than that of untreated organoids. While direct binding of topotecan and irinotecan with eGFP and eGFP-derivatives has not been reported, it is possible that they interact with these fluorescent proteins via other intracellular proteins to dampen fluorescence. Alternatively, it is possible topotecan and irinotecan have pleotropic effects, in which they affect the expression of other genes that indirectly regulate *UBE3A*.

## 5 Conclusion

Overall, this study develops a human-specific paternal *UBE3A* fluorescent reporter cell line. This model can be used for investigating the mechanism of UBE3A in driving AS phenotypes in neurodevelopmental cell types. Moreover, this reporter system could be a useful preclinical tool to screen therapeutic candidates targeting paternal *UBE3A* reactivation.

## 7 Author Contributions

GRK and SRS contributed equally to this work. GRK and SRS jointly decided, with agreement from all authors they all have the permission to list their names first on professional documents. Conceptualization: SRS, GRK, DS, AJK.

Methodology: SRS, GRK, DS, ZBY, LH, LL, JKH. Investigation: GRK, SRS, DS, ZBY, LH, LL, JKH, AJK.

Visualization: GRK, SRS, ZBY.

Resources: LL, JKH. Supervision: AJK.

Writing - Original draft: GRK, SRS, AJK.

Writing – Review and Editing: GRK, SRS, DS, ZBY, LH, LL, JKH, AJK

## 9 Acknowledgments

We thank Dr. Stormy Chamberlain (University of Connecticut Genetics and Genome Sciences, Farmington, CT) for graciously gifting the H9_*UBE3A m-/p-*_ cell line. We thank Daphne Collias and Dr. Chase Beisel for sharing a modified pX330 plasmid to generate pDS48. We thank Dr. Maria Theresa Fadri, Tyler J. Johnson, and Rachel Polak for their thoughtful insights and feedback.Figures 1A and S1A were created with BioRender.com.

## 12 Data Availability Statement

The datasets used to generate the graphs in this study can be found in the supplementary materials.

## References

Albrecht, Urs, James S Sutcliffe, Bruce M Cattanach, Colin V Beechey, Dawna Armstrong, Gregor Eichele, and Arthur L Beaudet. 1997. ‘Imprinted Expression of the Murine Angelman Syndrome Gene, Ube3a, in Hippocampal and Purkinje Neurons’. Nature Genetics 17 (1): 75– 78.

Almeida, Jacqueline Fátima Martins de, Ilaria Tonazzini, and Simona Daniele. 2025. ‘Molecular Aspects of Angelman Syndrome: Defining the New Path Forward’. Biomolecules and Biomedicine.

Angelman, Harry. 1965. ‘‘Puppet’Children a Report on Three Cases’. Developmental Medicine & Child Neurology 7 (6): 681–88.

Bazick, Hannah O, Hanqian Mao, Jesse K Niehaus, Justin M Wolter, and Mark J Zylka. 2024. ‘AAV Vector-Derived Elements Integrate into Cas9-Generated Double-Strand Breaks and Disrupt Gene Transcription’. Molecular Therapy 32 (11): 4122–37.

Burette, Alain C, Matthew C Judson, Susan Burette, Kristen D Phend, Benjamin D Philpot, and Richard J Weinberg. 2017. ‘Subcellular Organization of UBE3A in Neurons’. Journal of Comparative Neurology 525 (2): 233–51.

Burette, Alain C, Matthew C Judson, Alissa N Li, Edward F Chang, William W Seeley, Benjamin D Philpot, and Richard J Weinberg. 2018. ‘Subcellular Organization of UBE3A in Human Cerebral Cortex’. Molecular Autism 9:1–14.

Campbell, Benjamin C, Elisa M Nabel, Mitchell H Murdock, Cristina Lao-Peregrin, Pantelis Tsoulfas, Murray G Blackmore, Francis S Lee, Conor Liston, Hirofumi Morishita, and Gregory A Petsko. 2020. ‘mGreenLantern: A Bright Monomeric Fluorescent Protein with Rapid Expression and Cell Filling Properties for Neuronal Imaging’. Proceedings of the National Academy of Sciences 117 (48): 30710–21.

Chamberlain, Stormy J, Pin-Fang Chen, Khong Y Ng, Fany Bourgois-Rocha, Fouad Lemtiri-Chlieh, Eric S Levine, and Marc Lalande. 2010. ‘Induced Pluripotent Stem Cell Models of the Genomic Imprinting Disorders Angelman and Prader–Willi Syndromes’. Proceedings of the National Academy of Sciences 107 (41): 17668–73.

Chen, Pin-Fang. 2016. ‘Modeling Neurodevelopmental Disorders Involving Genomic Imprinting at Human Chromosome 15q11-Q13 Using iPSC and CRISPR/Cas9 Technology’. Doctoral Dissertation, University Connecticut.

Ciceri, Gabriele, Arianna Baggiolini, Hyein S Cho, Meghana Kshirsagar, Silvia Benito-Kwiecinski, Ryan M Walsh, Kelly A Aromolaran, et al. 2024. ‘An Epigenetic Barrier Sets the Timing of Human Neuronal Maturation’. Nature 626 (8000): 881–90.

Clarke, Maria T, Laura Remesal, Lea Lentz, Danielle J Tan, David Young, Slesha Thapa, Shalini R Namuduri, et al. 2024. ‘Prenatal Delivery of a Therapeutic Antisense Oligonucleotide Achieves Broad Biodistribution in the Brain and Ameliorates Angelman Syndrome Phenotype in Mice’. Molecular Therapy 32 (4): 935–51.

Condon, Kathryn H, Jianghai Ho, Camenzind G Robinson, Cyril Hanus, and Michael D Ehlers. 2013. ‘The Angelman Syndrome Protein Ube3a/E6AP Is Required for Golgi Acidification and Surface Protein Sialylation’. Journal of Neuroscience 33 (9): 3799–3814.

Dindot, Scott V, Barbara A Antalffy, Meenakshi B Bhattacharjee, and Arthur L Beaudet. 2008. ‘The Angelman Syndrome Ubiquitin Ligase Localizes to the Synapse and Nucleus, and Maternal Deficiency Results in Abnormal Dendritic Spine Morphology’. Human Molecular Genetics 17 (1): 111–18.

Dindot, Scott V, Sarah Christian, William J Murphy, Allyson Berent, Jennifer Panagoulias, Annalise Schlafer, Johnathan Ballard, et al. 2023. ‘An ASO Therapy for Angelman Syndrome That Targets an Evolutionarily Conserved Region at the Start of the UBE3A-AS Transcript’. Science Translational Medicine 15 (688): eabf4077.

Estridge, R Chris, Z Begum Yagci, Dilara Sen, Tyler J Johnson, Gautami R Kelkar, Travis S Ptacek, Jeremy M Simon, and Albert J Keung. 2025. ‘Loss of UBE3A Impacts Both Neuronal and Non-Neuronal Cells in Human Cerebral Organoids’. Communications Biology 8 (1): 1–19.

Filonova, Irina, Justin H Trotter, Jessica L Banko, and Edwin J Weeber. 2014. ‘Activity-Dependent Changes in MAPK Activation in the Angelman Syndrome Mouse Model’. Learning & Memory 21 (2): 98–104.

Fink, James J, Tiwanna M Robinson, Noelle D Germain, Carissa L Sirois, Kaitlyn A Bolduc, Amanda J Ward, Frank Rigo, Stormy J Chamberlain, and Eric S Levine. 2017. ‘Disrupted Neuronal Maturation in Angelman Syndrome-Derived Induced Pluripotent Stem Cells’. Nature Communications 8 (1): 15038.

Grier, Mark D, Robert P Carson, and Andre Hollis Lagrange. 2015. ‘Toward a Broader View of Ube3a in a Mouse Model of Angelman Syndrome: Expression in Brain, Spinal Cord, Sciatic Nerve and Glial Cells’. PloS One 10 (4): e0124649.

Han, Linna, Z Begum Yagci, and Albert J Keung. 2025. ‘A High Sensitivity Assay of UBE3A Ubiquitin Ligase Activity’. Methods.

Hillman, Paul R, Sarah GB Christian, Ryan Doan, Noah D Cohen, Kranti Konganti, Kory Douglas, Xu Wang, Paul B Samollow, and Scott V Dindot. 2017. ‘Genomic Imprinting Does Not Reduce the Dosage of UBE3A in Neurons’. Epigenetics & Chromatin 10:1–14.

Hsiao, Jack S, Noelle D Germain, Andrea Wilderman, Christopher Stoddard, Luke A Wojenski, Geno J Villafano, Leighton Core, Justin Cotney, and Stormy J Chamberlain. 2019. ‘A Bipartite Boundary Element Restricts UBE3A Imprinting to Mature Neurons’. Proceedings of the National Academy of Sciences 116 (6): 2181–86.

Huang, Hsien-Sung, John A Allen, Angela M Mabb, Ian F King, Jayalakshmi Miriyala, Bonnie Taylor-Blake, Noah Sciaky, et al. 2012. ‘Topoisomerase Inhibitors Unsilence the Dormant Allele of Ube3a in Neurons’. Nature 481 (7380): 185–89.

Jones, Kelly A, Ji Eun Han, Jason P DeBruyne, and Benjamin D Philpot. 2016. ‘Persistent Neuronal Ube3a Expression in the Suprachiasmatic Nucleus of Angelman Syndrome Model Mice’. Scientific Reports 6 (1): 28238.

Judson, Matthew C, Jason O Sosa-Pagan, Wilmer A Del Cid, Ji Eun Han, and Benjamin D Philpot. 2014. ‘Allelic Specificity of Ube3a Expression in the Mouse Brain during Postnatal Development’. Journal of Comparative Neurology 522 (8): 1874–96.

Kaphzan, Hanoch, Pepe Hernandez, Joo In Jung, Kiriana K Cowansage, Katrin Deinhardt, Moses V Chao, Ted Abel, and Eric Klann. 2012. ‘Reversal of Impaired Hippocampal Long-Term Potentiation and Contextual Fear Memory Deficits in Angelman Syndrome Model Mice by ErbB Inhibitors’. Biological Psychiatry 72 (3): 182–90.

Keute, Marius, Meghan T Miller, Michelle L Krishnan, Anjali Sadhwani, Stormy Chamberlain, Ronald L Thibert, Wen-Hann Tan, Lynne M Bird, and Joerg F Hipp. 2021. ‘Angelman Syndrome Genotypes Manifest Varying Degrees of Clinical Severity and Developmental Impairment’. Molecular Psychiatry 26 (7): 3625–33.

Khatri, Natasha, and Heng-Ye Man. 2019. ‘The Autism and Angelman Syndrome Protein Ube3A/E6AP: The Gene, E3 Ligase Ubiquitination Targets and Neurobiological Functions’. Frontiers in Molecular Neuroscience 12:109.

Kishino, Tatsuya, Marc Lalande, and Joseph Wagstaff. 1997. ‘UBE3A/E6-AP Mutations Cause Angelman Syndrome’. Nature Genetics 15 (1): 70–73.

Lancaster, Madeline A, Magdalena Renner, Carol-Anne Martin, Daniel Wenzel, Louise S Bicknell, Matthew E Hurles, Tessa Homfray, Josef M Penninger, Andrew P Jackson, and Juergen A Knoblich. 2013. ‘Cerebral Organoids Model Human Brain Development and Microcephaly’. Nature 501 (7467): 373–79.

LaSalle, Janine M, Lawrence T Reiter, and Stormy J Chamberlain. 2015. ‘Epigenetic Regulation of UBE3A and Roles in Human Neurodevelopmental Disorders’. Epigenomics 7 (7): 1213–28.

Lee, Dongwon, Wu Chen, Heet Naresh Kaku, Xinming Zhuo, Eugene S Chao, Armand Soriano, Allen Kuncheria, et al. 2023. ‘Antisense Oligonucleotide Therapy Rescues Disturbed Brain Rhythms and Sleep in Juvenile and Adult Mouse Models of Angelman Syndrome’. Elife 12:e81892.

Lee, Hyeong-Min, Ellen P Clark, M Bram Kuijer, Mark Cushman, Yves Pommier, and Benjamin D Philpot. 2018. ‘Characterization and Structure-Activity Relationships of Indenoisoquinoline-Derived Topoisomerase I Inhibitors in Unsilencing the Dormant Ube3a Gene Associated with Angelman Syndrome’. Molecular Autism 9:1–10.

Matsuura, Toshinobu, James S Sutcliffe, Ping Fang, Robert-Jan Galjaard, Yong-hui Jiang, Claudia S Benton, Johanna M Rommens, and Arthur L Beaudet. 1997. ‘De Novo Truncating Mutations in E6-AP Ubiquitin-Protein Ligase Gene (UBE3A) in Angelman Syndrome’. Nature Genetics 15 (1): 74–77.

McCoy, Eric S, Bonnie Taylor-Blake, Megumi Aita, Jeremy M Simon, Benjamin D Philpot, and Mark J Zylka. 2017. ‘Enhanced Nociception in Angelman Syndrome Model Mice’. Journal of Neuroscience 37 (42): 10230–39.

Meng, Linyan, Amanda J Ward, Seung Chun, C Frank Bennett, Arthur L Beaudet, and Frank Rigo. 2015. ‘Towards a Therapy for Angelman Syndrome by Targeting a Long Non-Coding RNA’. Nature 518 (7539): 409–12.

Munshi, Shashini. 2019. ‘Modeling Human Brain Diseases Using Pluripotent Stem Cells’. Doctoral Dissertation, Erasmus University Rotterdam.

Rougeulle, Claire, Heather Glatt, and Marc Lalande. 1997. ‘The Angelman Syndrome Candidate Gene, UBE3AIE6-AP, Is Imprinted in Brain’. Nature Genetics 17 (1): 14–15.

Schindelin, Johannes, Ignacio Arganda-Carreras, Erwin Frise, Verena Kaynig, Mark Longair, Tobias Pietzsch, Stephan Preibisch, et al. 2012. ‘Fiji: An Open-Source Platform for Biological-Image Analysis’. Nature Methods 9 (7): 676–82.

Schmid, Ralf S, Xuefeng Deng, Priyalakshmi Panikker, Msema Msackyi, Camilo Breton, James M Wilson, and others. 2021. ‘CRISPR/Cas9 Directed to the Ube3a Antisense Transcript Improves Angelman Syndrome Phenotype in Mice’. The Journal of Clinical Investigation 131 (5).

Sen, Dilara, Zuzana Drobna, and Albert J Keung. 2021. ‘Evaluation of UBE3A Antibodies in Mice and Human Cerebral Organoids’. Scientific Reports 11 (1): 6323.

Sen, Dilara, Alexis Voulgaropoulos, Zuzana Drobna, and Albert J Keung. 2020. ‘Human Cerebral Organoids Reveal Early Spatiotemporal Dynamics and Pharmacological Responses of UBE3A’. Stem Cell Reports 15 (4): 845–54.

Sirois, Carissa. 2018. ‘Generation of Isogenic Human Pluripotent Stem Cell-Derived Neurons to Establish a Molecular Angelman Syndrome Phenotype and to Study the UBE3A Protein Isoforms’. Doctoral Dissertation, University Connecticut.

Sirois, Carissa L, Judy E Bloom, James J Fink, Dea Gorka, Steffen Keller, Noelle D Germain, Eric S Levine, and Stormy J Chamberlain. 2020. ‘Abundance and Localization of Human UBE3A Protein Isoforms’. Human Molecular Genetics 29 (18): 3021–31.

Sonzogni, Monica, Peipei Zhai, Edwin J Mientjes, Geeske M van Woerden, and Ype Elgersma. 2020. ‘Assessing the Requirements of Prenatal UBE3A Expression for Rescue of Behavioral Phenotypes in a Mouse Model for Angelman Syndrome’. Molecular Autism 11:1–12.

Stanurova, Jana, Anika Neureiter, Michaela Hiber, Hannah de Oliveira Kessler, Kristin Stolp, Roman Goetzke, Diana Klein, Agnes Bankfalvi, Hannes Klump, and Laura Steenpass. 2016. ‘Angelman Syndrome-Derived Neurons Display Late Onset of Paternal UBE3A Silencing’. Scientific Reports 6 (1): 30792.

Sutcliffe, James S, Yong-hui Jiang, Robert-Jan Galjaard, Toshinobu Matsuura, Ping Fang, Takeo Kubota, Susan L Christian, et al. 1997. ‘The E6–AP Ubiquitin–Protein Ligase (UBE3A) Gene Is Localized within a Narrowed Angelman Syndrome Critical Region’. Genome Research 7 (4): 368–77.

Vihma, Hanna, Kelin Li, Anna Welton-Arndt, Audrey L Smith, Kiran R Bettadapur, Rachel B Gilmore, Eric Gao, et al. 2024. ‘Ube3a Unsilencer for the Potential Treatment of Angelman Syndrome’. Nature Communications 15 (1): 5558.

Vu, Thanh H, and Andrew R Hoffman. 1997. ‘Imprinting of the Angelman Syndrome Gene, UBE3A, Is Restricted to Brain’. Nature Genetics 17 (1): 12–13.

Wolter, Justin M, Hanqian Mao, Giulia Fragola, Jeremy M Simon, James L Krantz, Hannah O Bazick, Baris Oztemiz, Jason L Stein, and Mark J Zylka. 2020. ‘Cas9 Gene Therapy for Angelman Syndrome Traps Ube3a-ATS Long Non-Coding RNA’. Nature 587 (7833): 281– 84.

Zampeta, F Isabella, Monica Sonzogni, Eva Niggl, Bas Lendemeijer, Hilde Smeenk, Femke MS de Vrij, Steven A Kushner, Ben Distel, and Ype Elgersma. 2020. ‘Conserved UBE3A Subcellular Distribution between Human and Mice Is Facilitated by Non-Homologous Isoforms’. Human Molecular Genetics 29 (18): 3032–43.

