## Supplementary Material for "A Human Angelman Syndrome Class II Pluripotent Stem Cell line with Fluorescent Paternal *UBE3A* Reporter"

### 1 Supplementary Figures and Tables

#### 1.1 Supplementary Figures

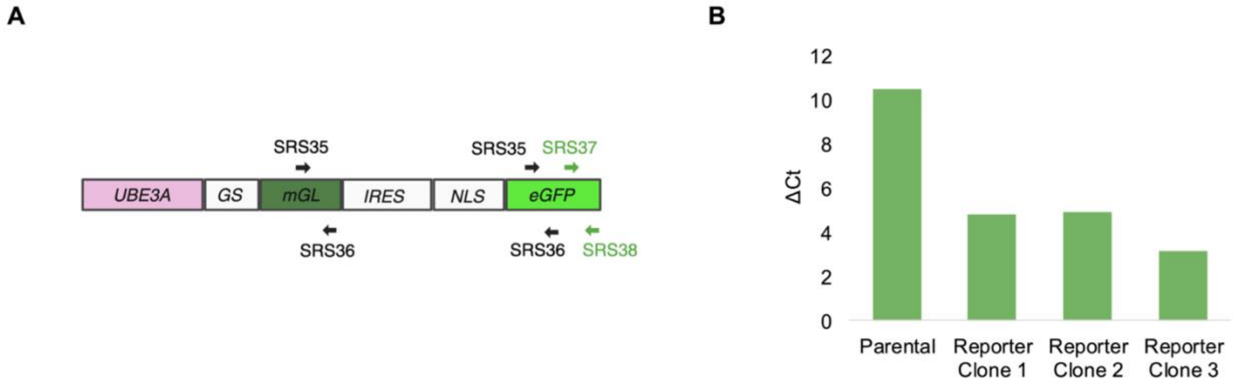


**Figure S1.** qPCR measurements to confirm reporter construct expression. A) Primer target regions within the transcript. Data from primers SRS35 and SRS36 (black) is shown in Figure 1C. B) Data for region amplified by primers SRS37 and SRS38. ΔCt values relative to GAPDH presented for the parental iPSCs and the three monoclonal edited reporter iPSCs.

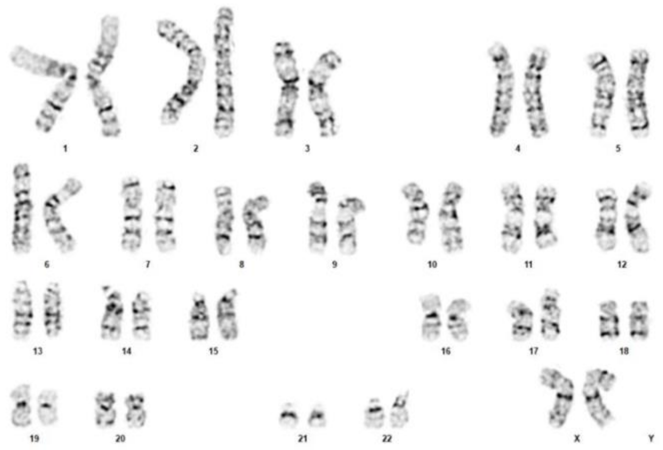

**Figure S2.** AS Class II deletion parental cell line shows normal karyotype 46 XX. The karyotype analysis was performed by LabCorp.

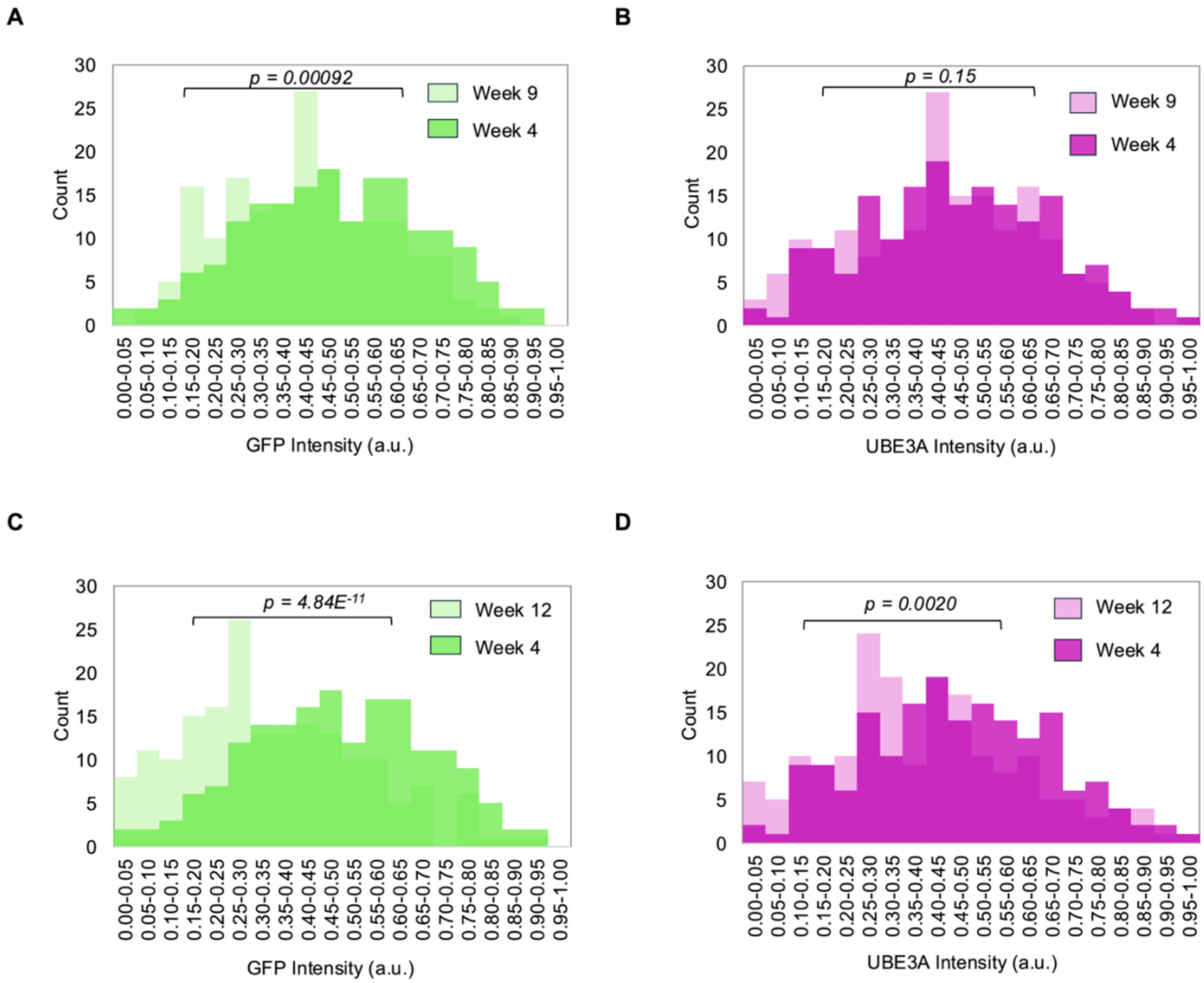


**Figure S3.** Intensity histograms from confocal images for nuclei in TUJ1+ cells, comparing A) GFP and B) UBE3A at week 4 to week 9, and C) GFP and D) UBE3A at week 4 to week 12. Populations were compared using a two-sample t-test, n=180 cells (3 organoids). a.u. – arbitrary units.

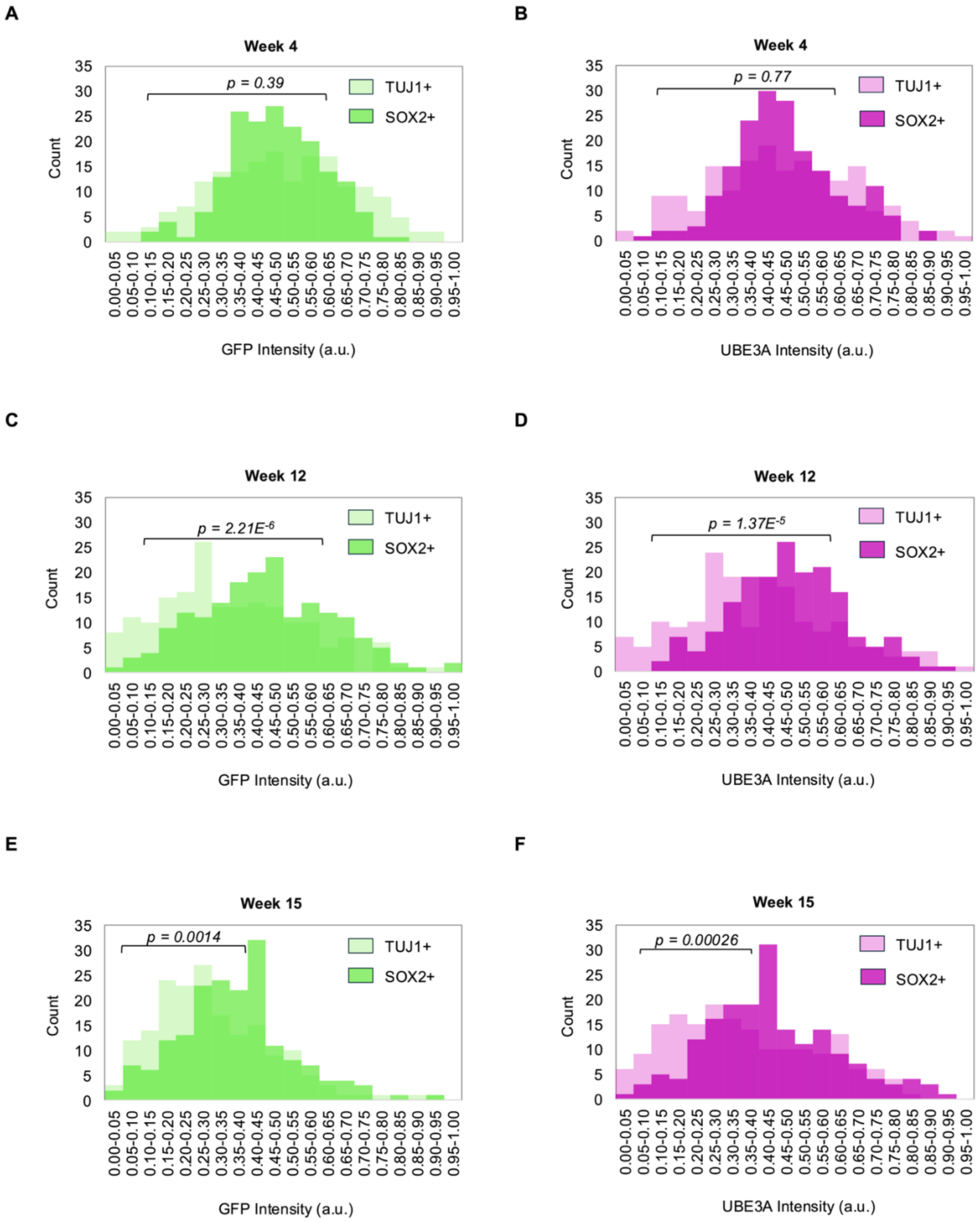
A
Week 4
p = 0.39
TUJ1+
SOX2+
Count
GFP Intensity (a.u.)
B
Week 4
p = 0.77
TUJ1+
SOX2+
Count
UBE3A Intensity (a.u.)
C
Week 12
p = 2.21E-6
TUJ1+
SOX2+
Count
GFP Intensity (a.u.)
D
Week 12
p = 1.37E-5
TUJ1+
SOX2+
Count
UBE3A Intensity (a.u.)
E
Week 15
p = 0.0014
TUJ1+
SOX2+
Count
GFP Intensity (a.u.)
F
Week 15
p = 0.00026
TUJ1+
SOX2+
Count
UBE3A Intensity (a.u.)

**Figure S4.** GFP and UBE3A intensity histograms from confocal images for nuclei in TUJ1+ cells compared to SOX2+ nuclei at A) and B) week 4, C) and D) week 12, and E) and F) week 15. Populations were compared using a two-sample t-test, n = 180 cells (3 organoids). a.u.- arbitrary units.

### 1.2 Supplementary Tables

**Table S1:** Primer Sequences.

| Primer Name | Experiment | Sequence - 5' to 3' |
|---|---|---|
| SRS21 | Genomic PCR Screening – Round 1 and 2 | ACCTTGCATTCCTCGTCACA |
| SRS24 | Genomic PCR Screening – Round 1 | CCTCACATTGCCAAAAGACG |
| DSP206 | Genomic PCR Screening – Round 2 | CACTGATCACGTGCCTCGATA |
| SRS35 | qPCR: mGL-eGFP | AAGCAGAAGAACGGCATCAA |
| SRS36 | qPCR: mGL-eGFP | GGGGGTGTTCTGCTGGTAGT |
| SRS37 | qPCR: eGFP only | AGTCCGCCCTGAGCAAAGA |
| SRS38 | qPCR: eGFP only | TCCAGCAGGACCATGTGATC |

**Table S2:** Green fluorescence intensity (a.u.) data from flow cytometry experiments.

| Sample | %GF+ Cells | For GF+ population | | |
|---|---|---|---|---|
| | | Geometric Mean | Median | Mode |
| Reporter Organoids + Vehicle Control | 23.2 | 2.88 | 2.65 | 2.02 |
| Reporter Organoids + 1μM Topotecan | 23.8 | 2.93 | 2.74 | 2.13 |
| Reporter Organoids + 1μM Irinotecan | 28.8 | 3.01 | 2.82 | 2.07 |
| Reporter iPSCs (Positive Control) | 48.4 | 3.04 | 2.97 | 2.86 |
| Parental iPSCs (Negative Control) | 0.14 | 2.68 | 2.36 | 2.02 |
